# SgTiler: a fast method to design tiling sgRNAs for CRISPR/Cas9 mediated screening

**DOI:** 10.1101/217166

**Authors:** Musaddeque Ahmed, Housheng Hansen He

## Abstract

**Summary:** Screening of genomic regions of interest using CRISPR/Cas9 is getting increasingly popular. The system requires designing of single guide RNAs (sgRNAs) that can efficiently guide the Cas9 endonuclease to the targeted region with minimal off-target effects. Tiling sgRNAs is the most effective way to perturb regulatory regions, such as promoters and enhancers. sgTiler is the first tool that provides a fast method for designing tiling sgRNAs.

**Availability and Implementation:** sgTiler is a command line tool that requires only one command to execute. Its source code is freely available on the web at https://github.com/HansenHeLab/sgTiler. sgTiler is implemented in Python and supported on any platform with Python and Bowtie.

## Introduction

The Clustered Regularly Interspaced Short Palindromic Repeats (CRISPR) system consists of two parts – a non-specific endonuclease Cas9 and a short guide RNA (sgRNA). The sgRNA is a synthetic RNA consists of a scaffold sequence necessary for Cas9 function and a ~20nt spacer sequence that can recognize genomic region. The spacer sequence guides the Cas9 to bind to targeted region where the endonuclease creates a double-stranded cleavage (Le Cong et al., 2013; Mali et al., 2013). This makes CRISPR/Cas9 system a highly preferable method to mutate/inactivate any region for both its efficiency and specificity. The simplicity of the method has attracted many screening studies in both cell culture and whole organisms and the number of screening studies is increasing rapidly (Korkmaz et al., 2016; Shalem et al., 2014; Shi et al., 2015; Wang et al., 2014; Zhu et al., 2016). The widespread popularity of CRISPR/Cas9-mediated screening also led to development of many computational tools to predict sgRNAs (reviewed in (Chuai, Wang, & Liu, 2016)). However, most of the currently available tools are very slow and online-only, restricting it’s use for larger number of targets; and none of the tools is intended for tiling sgRNAs over the input regions. Tiling sgRNAs are overlapping sgRNAs maximizing the possibility of perturbing the target region, provided an effective approach for targeting regulatory regions. Here we present sgTiler that provides a fastest and efficient method to design tiling sgRNAs with novel approach for off-target effect calculation and optimization of number of sgRNAs.

## Methods

The tiling sgRNAs are designed in two major steps – first, identify candidate sgRNAs within the input region and second, optimize the distribution of sgRNAs. SgTiler identifies any sequence of the length of the spacer that meets the following critiera: 1) ends with the PAM, 2) meets certain GC content requirement, and 3) does not contain a poly T sequence. This is important because sgRNAs with too low or too high GC content or UUUU/TTTT sequences have been found to have lower cleavage efficiency (Gilbert et al., 2014; T. Wang et al., 2014). Then sgTiler calculates the efficiency of each sgRNA based on the weight matrix of each position as previously described (Doench et al., 2014; Heigwer et al., 2016). sgTiler retains any sgRNA that 1) maps uniquely within the input region, 2) does not map anywhere else in the genome with less than 2 mismatches and 3) it’s efficiency score can be calculated (Figure 1). In the next step, sgTiler implements a novel approach to calculate the off-target potential (OTP) of each candidate sgRNA by utilizing user-provided annotations of important genomic features. For example, for *n* number of genomic features, OTP score of a sgRNA would be 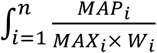 is the number of intersection of the given sgRNA with the *i*th feature, *MAX* is the maximum intersection of any sgRNA within the same input region and *W* is a weight given to the feature. This relative scoring approach contributes to the inclusive nature of the tool. In its default setting, sgTiler requires two annotation files –exome coordinates and coordinates of open chromatin (DNaseI hypersensitive sites or histone marks). sgTiler weighs the exome regions twice as high as the open chromatin, and in such a way that the worst OTP score is 1. It is particularly important that, even with mismatches, spacer sequence should not be complimentary to any genic or regulatory region. Thus, an OTP score approaching zero is preferable. This ensures that even if a spacer sequence maps to elsewhere in the genome, it will be less likely to cause any effect minimizing the false discovery rate.

**Fig 1:**
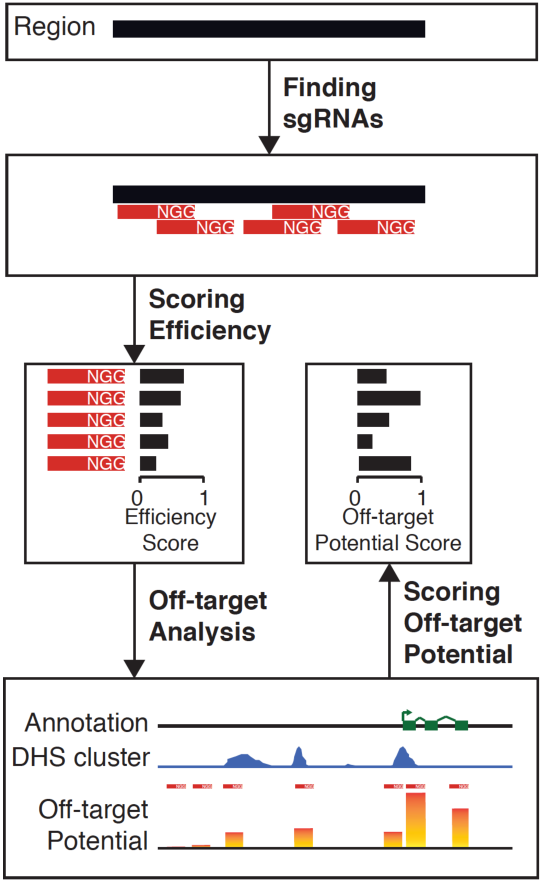
Schematic for identification of candidate sgRNAs. sgRNAs with an efficiency score close to 1 is predicted to have high cleavage efficiency, and with Off-target Potential score of 0 has least likelihood for off-target effect.

In the second step of tiling-sgRNA design, sgTiler implements an optimization technique to reduce the number of overlapping sgRNAs. One of the key factors in designing tiling sgRNAs for regulatory elements (e.g., promoters or enhancers) is the coverage of the entire region. It is important to distribute the sgRNAs in a way that they span across the whole region without leaving large gaps in between. sgTiler optimizes the number and distribution of sgRNAs in three sequential steps. For any particular input region, sgTiler first determines evenly distributed nucleotide positions (termed as “even points”) using Bresenham’s line algorithm. Next, the algorithm selects the sgRNAs around the even points within a certain nucleotide window. For even points surrounded by more than one sgRNAs, sgTiler picks the sgRNA by prioritizing in the following order: lower off-target potential, higher efficiency and closeness to the even point. In the third step, the algorithm optimizes the design by loosening the threshold for the even points devoid of sgRNA. This ensures that the sgRNAs span the most nucleotides in the input region, as well as that the sgRNAs are not sporadically clustered. The optimization typically reduces the number of sgRNAs by ~50% per region without decreasing the coverage, consequently making room for more regions in library preparation.

## Results

SgTiler is written in Python. It requires bowtie and one simple command line to execute. SgTiler requires three input files – the input fasta file, a bed file with exome coordinates and one bed file of regulatory regions. The regulatory regions can be typically active histone marks or DNaseI hypersensitive sites. The tool output four text files – list of all candidate sgRNAs, list of filtered sgRNAs, list of sgRNA details for the input regions and a summary report with important statistics. In addition, sgTiler also produces two graphical summaries of sgRNAs and sequence coverage per input region. SgTiler also generates graphical representation of distribution of sgRNA for each individual input region. Combining the overall statistics, user can predict the success of the screening.

To the best of our knowledge, sgTiler is the first tool that enables one to quickly design tiling sgRNAs for any input DNA sequence, including genes or regulatory regions. It can also be used for identifying sgRNAs in genic regions, not tiling, by turning off the optimization step. SgTiler is significantly faster than the tools that are currently available for designing sgRNAs, even though none of the currently available tools are designed for tiling. Furthermore, the optimization step disperses the sgRNAs to optimally cover the entire region with minimum sgRNAs. This enables the maximum use of the library and makes room for more regions.

